# Alterations in the gut microbiome of whiteleg shrimp (*Penaeus vannamei*) postlarvae following exposure to an AHPND-causing strain of *Vibrio parahaemolyticus*

**DOI:** 10.1101/2023.07.15.548467

**Authors:** Manuel J. Beltrán, Juan Quimi Mujica, Benoit Diringer, Sergio P. Barahona

## Abstract

Acute Hepatopancreatic Necrosis Disease (AHPND), attributed to the production of PirA/PirB toxins by certain Vibrio parahaemolyticus strains, poses a significant threat to global shrimp aquaculture, causing substantial mortality and economic losses. To enhance our understanding of this disease within a closed culture system on the northern coast of Peru, we conducted a comparative analysis of the gut microbiomes between healthy and diseased postlarvae. Diseased postlarvae were obtained through exposure to an AHPND-causing strain of V. parahaemolyticus. Five healthy and five diseased postlarvae were randomly sampled from experimental rearing tanks, and their medial guts were extracted. High-throughput sequencing targeting the V4 region of the 16S rRNA gene was employed for amplicon library construction, and assessments of alpha and beta diversities, as well as taxonomic composition, were conducted. Our results revealed reduced diversity and distinct compositional profiles in the gut microbiomes of diseased postlarvae. The order Rhodobacteriales was dominant in the gut microbiomes of healthy postlarvae, while the order Vibrionales (including an unassigned genus within Vibrionales, Vibrio, and Pseudoalteromonas) exhibited the highest abundance in diseased postlarvae. In conclusion, exposure to an AHPND-causing strain of V. parahaemolyticus induces significant dysbiosis in the gut microbiome of whiteleg shrimp postlarvae.

## INTRODUCTION

Whiteleg shrimp *Penaeus vannamei* (Boone, 1931) stands as the most extensively farmed crustacean worldwide (FAO 2022). However, the intensification of shrimp aquaculture has been accompanied by an increased incidence of infectious outbreaks caused by various pathogens, including bacteria, viruses, protozoa, and fungi (Gunalan et al. 2014). Among these, Acute Hepatopancreatic Necrosis Disease (AHPND), commonly referred to as Early Mortality Syndrome (EMS), stands out as the most severe bacterial affliction affecting farmed shrimp populations (Chen et al. 2017). The detrimental impact of AHPND on the shrimp farming industry has been well-documented, resulting in substantial economic losses (Stentiford et al. 2012, Thitamadee et al. 2016, Holt et al. 2020). This disease primarily targets postlarvae and juveniles, leading to complete mortality within days of the initial manifestation of signs (Soto-Rodriguez et al. 2015, Restrepo et al. 2018). The first recorded outbreak of AHPND occurred in Hainan, China, in 2009, subsequently spreading to shrimp farms across Asia and the Americas (Soto-Rodriguez et al. 2015, Chen et al. 2017). The etiological agents responsible underlying AHPND encompass *Vibrio parahaemolyticus* strains carrying a ∼70 Kb plasmid housing the toxin-encoding genes PirA and PirB (Lee et al. 2015). Nevertheless, it is worth highlighting that other *Vibrio* species with the ability to generate AHPND have also been documented (Liu et al. 2018, Restrepo et al. 2018).

The intricate relationship between a host organism and its resident gut microbiome has garnered substantial interest within the scientific community (Holt et al. 2020). Shrimps, being one of the key players in aquaculture, heavily rely on a balanced gut microbiome for crucial physiological processes such as nutrient digestion, immune modulation, and growth regulation (Garibay-Valdez et al. 2020). Several studies conducted in shrimp species have highlighted the remarkable sensitivity of gut microbiomes to various factors, including infectious diseases, environmental stressors, developmental stages, dietary changes, and culture conditions (Zhang et al. 2014, Xiong et al. 2015, Huang et al. 2016, Zhu et al. 2016, Landsman et al. 2019, Liu et al. 2019, Pei et al. 2019). Leveraging high-throughput sequencing techniques targeting the V4 hypervariable region of the 16S rRNA gene, which serves as the gold-standard phylogenetic marker for bacterial community analysis (Yang et al. 2016), has revolutionized our understanding of the structure and functionality of shrimp gut microbiomes (Holt et al. 2020). Consequently, microbiome profiling has emerged as a promising biomarker for assessing the presence and impact of infectious diseases in shrimp aquaculture (Xiong et al. 2015, Dai et al. 2018, Holt et al. 2020).

Several studies have examined the gut microbiome dynamics in relation to AHPND in different geographic regions, such as Ecuador, Mexico, and China (Chen et al. 2017, Liu et al. 2018, Afrin et al. 2022, Reyes et al. 2022). However, studies focusing on AHPND-associated gut microbiomes in Peru are currently limited (Intriago et al. 2018). Considering that the gut microbiome composition can also be influenced by ecological roles and geographical factors (Bass et al. 2019), it is imperative to expand our understanding through further investigations. Thus, the present study aims to characterize the alterations in diversity and composition within the gut microbiota of whiteleg shrimp postlarvae upon exposure to an AHPND-causing *V. parahaemolyticus* strain. By comparing healthy and diseased individuals within a closed culture system located on the northern coast of Peru, we seek to provide valuable insights into the microbial changes associated with AHPND in this region.

## MATERIALS AND METHODS

The experiment started in August 2021 at Centro Colectivo Educativo Experimental de Biología y Biotecnología Acuática y Acuícola de Puerto Pizarro (CEBAP) in Tumbes, located on the northern coast of Peru. The rearing process adhered to standardized conditions, wherein a 1100-L tank was utilized, and weekly water changes were conducted using nearby mangrove water sources. The water temperature ranged from 22 to 31°C, salinity maintained at 10, pH levels ranged between 7.5 and 8.5, ammonium concentration ranged from 0 to 4 ppt, nitrite levels were below 0.5, and nitrate concentrations ranged from 0 to 10 ppt. A total of 500 postlarvae measuring 2 cm in length and weighing 4 g were randomly allocated and evenly distributed among ten 50-L tanks, with each tank accommodating 50 postlarvae. Five tanks were assigned to the “healthy” group, while the remaining five tanks comprised the “diseased” group. The postlarvae in the “healthy” tanks were reared under standard conditions for a duration of four weeks. In contrast, the postlarvae in the “diseased” tanks were subjected to an immersion method using a PirA/PirB toxin-producing strain of *V. parahaemolyticus* to simulate a genuine infection, as previously described (Lee et al. 2015, Liu et al. 2018). Subsequently, signs of disease and significant mortality (>90%) were observed within the diseased postlarvae tanks. Following the four-week rearing period, a random sampling approach was employed, selecting a single individual from each tank, resulting in five healthy postlarvae (1 to 5-BD) and five surviving diseased postlarvae (6 to 10-BD). Sacrificial procedures were performed by subjecting the ten selected shrimps to a heat shock protocol (−1ºC min^-1^) (Weineck et al. 2018, Diggles 2019) within the facilities of Incabiotec S.A.C. The medial guts were promptly extracted by making an incision along the midline of the exoskeleton utilizing aseptic techniques to minimize external microbial contamination.

DNA extractions were performed using the ZymoBIOMICS™ DNA Miniprep Kit (Zymo Research, USA) according to the manufacturer’s instructions. Subsequently, DNA concentrations and purities were measured with NanoDrop™ Spectophotometer (Thermo Scientific™, USA). Genomic DNA isolates were stored at -20ºC until further genomic library preparation. The Earth Microbiome Project Illumina Protocol (https://earthmicrobiome.org/protocols-and-standards/16s/) was employed, along with the primer-pair 515FB/860R (Appril et al. 2015, Parada et al. 2016) to amplify and sequence the V4 hypervariable region of the 16S rRNA gene. A paired-end 2×200 bp MiSeq run was conducted. Individual-specific barcodes were incorporated for a further efficient demultiplexing (Suppl. Table 1).

The analysis of 16S rRNA gene sequences was performed using the command-line interface of QIIME 2 (Bolyen et al. 2019), following the recommended bioinformatic pipeline. Raw fastq sequences were imported into QIIME 2 as “qza” artifacts, along with the corresponding metadata file (Table S1). Non-biological sequences such as barcodes, Illumina adapters, and primers were eliminated, and demultiplexing was carried out using the “Cutadapt” plugin within QIIME 2 (Martin, 2011). Denoising, removal of singletons, chimera removal, trimming of low-quality bases (Phred score <33), forward/reverse concatenation, and dereplication (sequence clustering at 100% similarity) were performed using the “q2-dada2” plugin in QIIME 2 (Callahan et al. 2016). This process generated a table of Amplicon Sequence Variants (ASVs) or “representative sequences”.

Alpha diversity, including ASV richness and three diversity indexes (Shannon, Simpson, and Pielou), was calculated for each microbiome using QIIME 2. The non-parametric Mann-Whitney U test (also known as Wilcoxon rank-sum test) in R software (R Core Team 2013) was employed to assess significant differences in alpha diversities between the two treatments (healthy *vs*. diseased). Jitter violin plots, comparing the treatments, were performed using the R package ggplot2 (Wickham 2016). To ensure adequate sampling, rarefaction curves based on ASV richness and Shannon index were generated in QIIME 2. For assessing differences in beta diversity between treatments, Principal Coordinate Analysis (PCoA) plots were constructed based on Bray-Curtis dissimilarity, Jaccard, weighted Unifrac, and unweighted Unifrac (Lozupone et al. 2005). The significance of dissimilarities was determined using non-parametric PERMANOVA tests with 999 permutations in QIIME 2.

Taxonomic assignments were performed on the ASVs using the “classify-sklearn” (q2sk) method, which is available in the QIIME 2’s “q2-feature-classifier” plugin (Bokulich et al. 2018). A pre-trained Naive Bayes taxonomy classifier, specifically trained for the “Greengenes2 2022.10 from 515F/806R region of sequences” database (McDonald et al. 2012), was utilized. This classifier is publicly available at https://docs.qiime2.org/2023.5/data-resources. ASV relative abundance data was obtained using the “collapse” method within the QIIME 2’s “taxa” plugin. Barplots were generated to illustrate the taxonomic categories, and a heatmap was constructed to visualize differential abundances at the phylum level.

## RESULTS

An average of 34,751 reads per gut microbiome of shrimp postlarvae was obtained. After quality control using the QIIME 2’s “q2-dada2” plugin, 76.03% of the sequences passed, resulting in an average of 26,430 reads (Table S2). A total of 293 ASVs with an average length of 253 bp were identified. The most abundant ASV accounted for 64,177 sequences, while the least frequent ASV was represented by only two sequences. Alpha diversity measurements (Table S1) and violin plots comparing alpha diversities between treatments using the Mann-Whitney U test (Fig. 1) are presented. The diseased postlarvae’s gut microbiomes exhibited significantly lower diversity compared to the healthy ones, as indicated by the Shannon, Simpson, and Pielou indexes (Table S3, Fig. 1). Rarefaction curves demonstrated that the sequencing effort was sufficient to capture the species richness of the microbiomes (Fig. 2). Beta diversity PCoA plots revealed distinct differences (PERMANOVA, *P* < 0.05) between the microbiomes of the two treatments (Figs. 3a-b). Weighted and unweighted UniFrac analysis, which incorporates phylogenetic information, further highlighted the dissimilar phylogenetic relationships between the two treatments (Figs. 3c-d). Notably, the microbiome of diseased postlarvae 10-BD exhibited remarkable divergence from the other samples, a finding that will be further discussed in subsequent sections.

**Figure 1.**
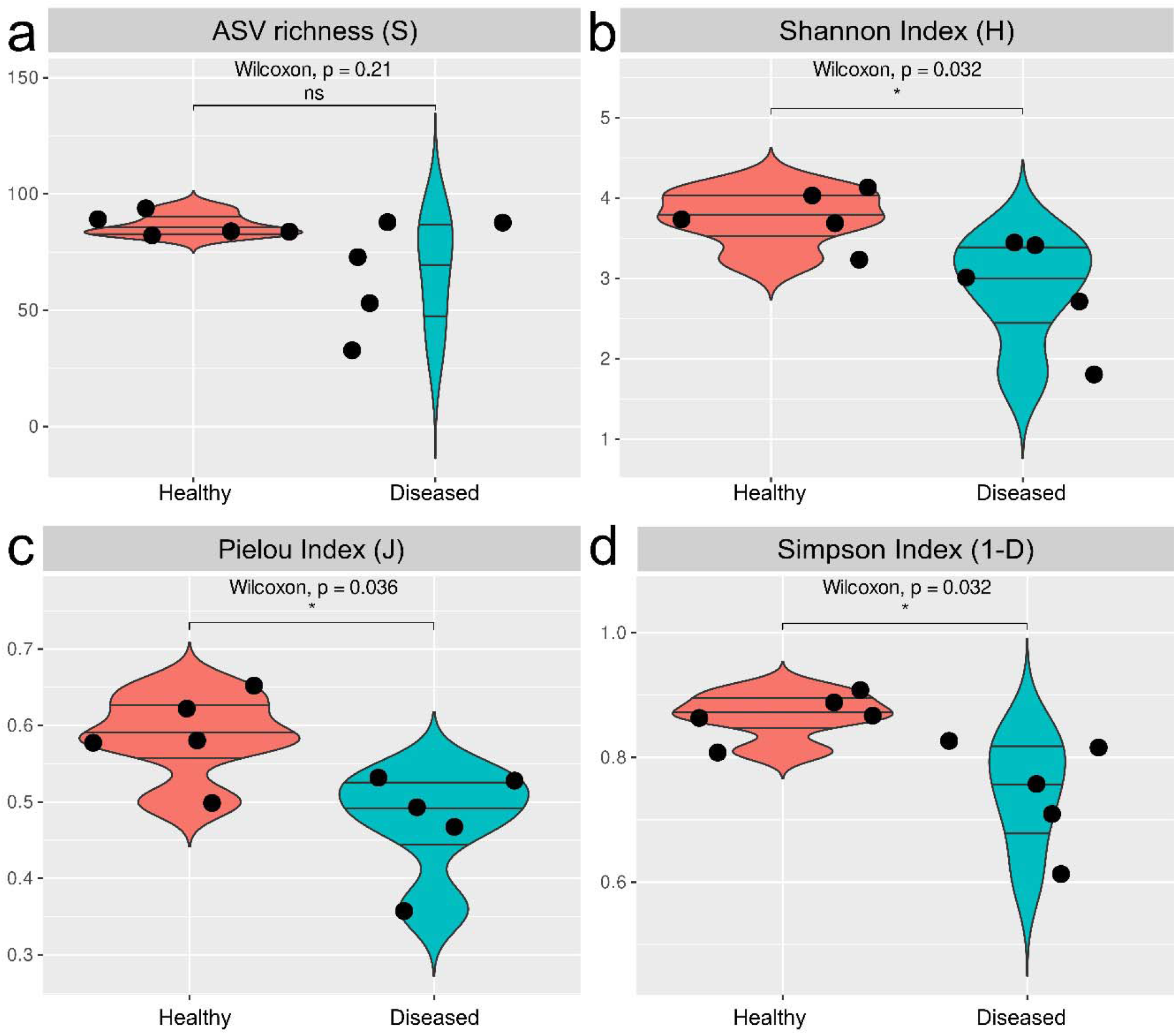
Comparison of alpha diversities between healthy and diseased postlarvae’s gut microbiomes shown as jitter violin plots. Measurements included (a) ASV richness, (b) Shannon Index, (c) Pielou Evenness Index and (d) Simpson Index. Statistical significances (p-value) of the Mann-Whitney U tests are indicated (ns = non-significant, * p < 0.05)

**Figure 2.**
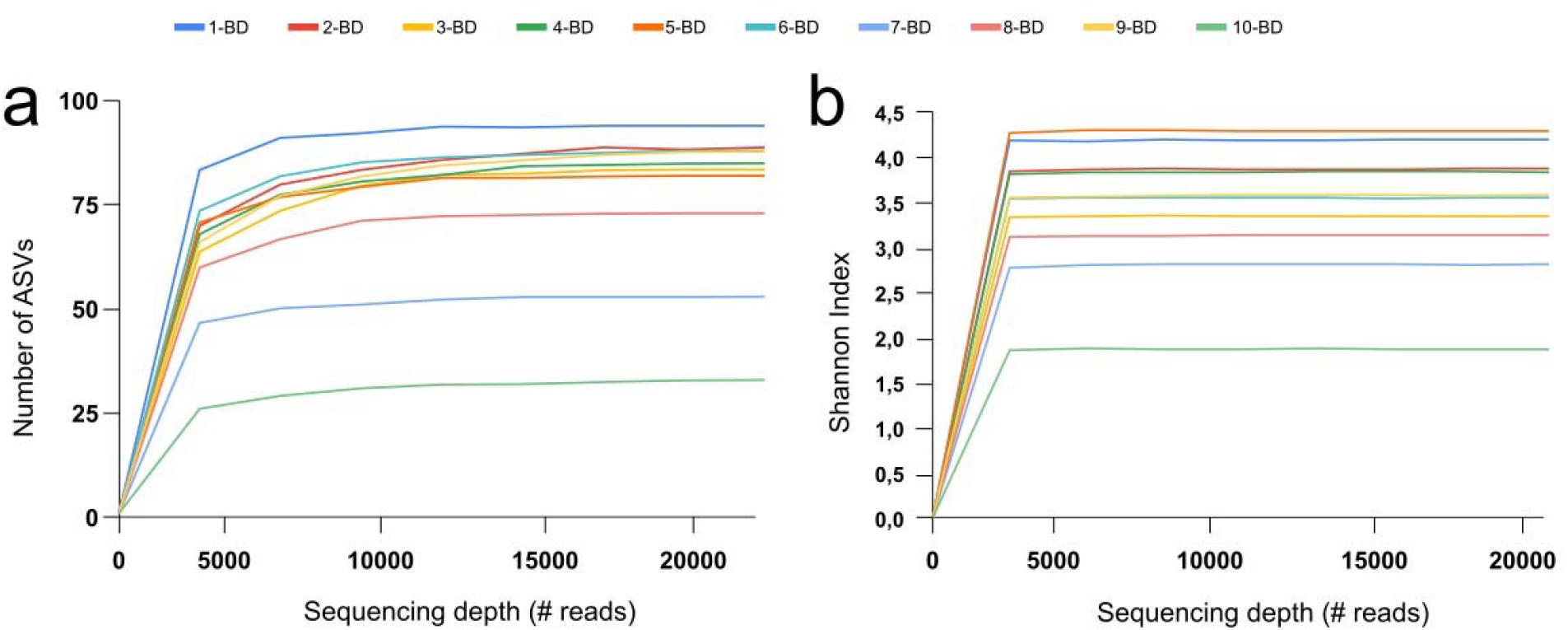
Rarefaction plots based on (a) ASV richness and (b) Shannon index

**Figure 3.**
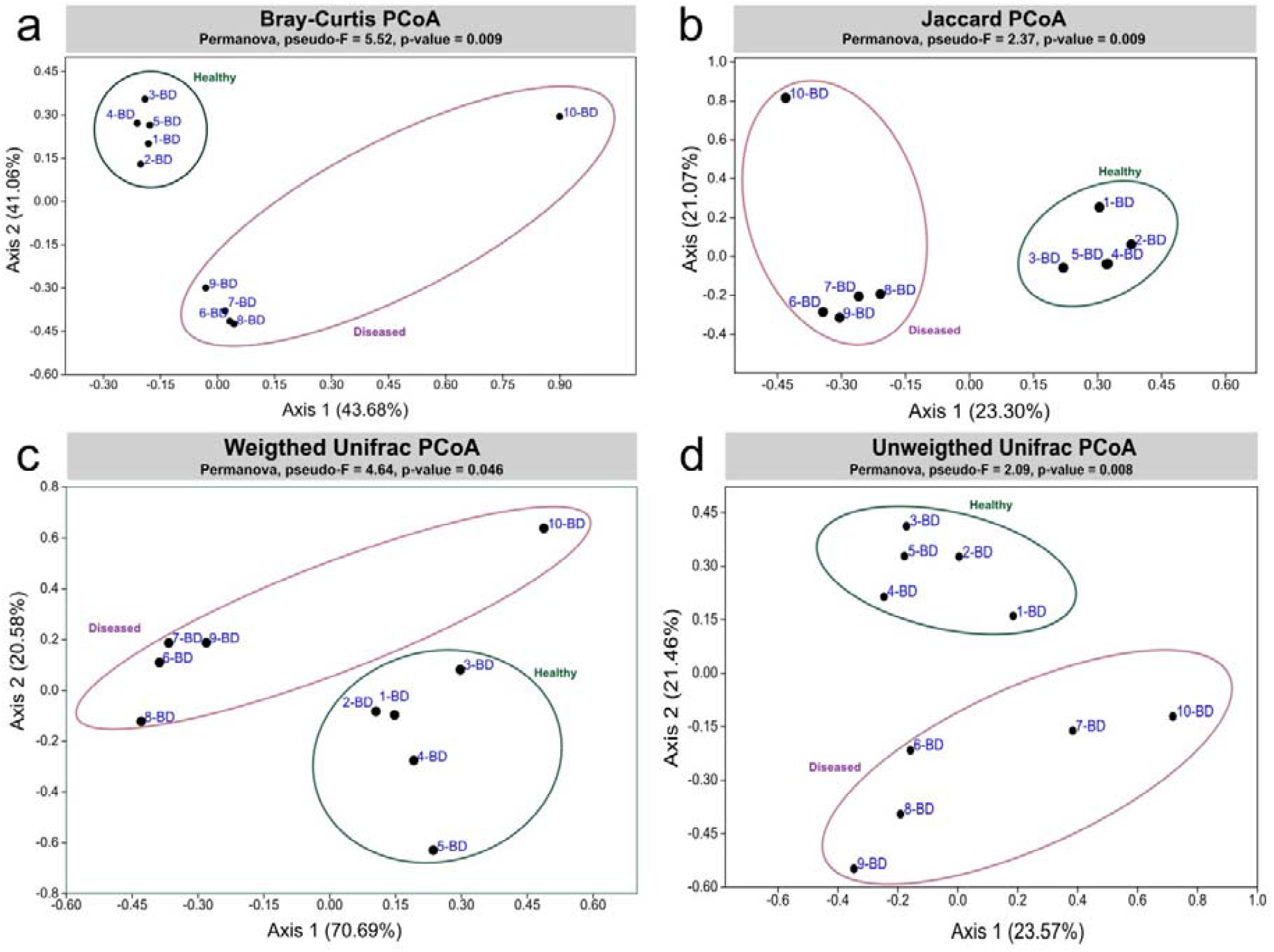
Beta diversity: Principal Coordinates Analysis (PCoA) plots based on (a) Bray-Curtis dissimilarity, (b) Jaccard Index, (d) Weighted Unifrac and (d) Unweighted Unifrac.>Explained variance of each axis are indicated as percentages. Healthy and diseased postlarvae’s gut microbiomes are indicated as dark dots and clustered into green and dark-pink ellipses respectively.

Proteobacteria emerged as the predominant phylum inhabiting the gut microbiomes of healthy postlarvae. In contrast, the gut microbiomes of diseased postlarvae exhibited a near-exclusive dominance of Proteobacteria compared to the healthy counterparts (Figs. 4a-b, 6, Table S4). Bacteroidetes ranked as the second most abundant phylum in healthy postlarvae’s gut microbiomes, with relative abundances of 18.7 and 29.43% observed in healthy postlarvae 4 and 5-BD, respectively. The diseased postlarvae’s gut microbiomes demonstrated reduced abundance of Bacteroidetes. Notably, the gut microbiome of postlarvae 1-BD exhibited a substantial presence of the phylum Actinobacteria (21.39%). In healthy postlarvae’s gut microbiomes, the class Alphaproteobacteria surpassed Gammaproteobacteria in abundance, whereas the diseased counterparts prominently displayed higher levels of Gammaproteobacteria, excluding postlarvae 10-BD. Class Cytophagia exhibited greater prevalence in healthy shrimp compared to the diseased ones (Fig. 5a). The order Rhodobacterales predominated in the gut microbiomes of healthy shrimp postlarvae, while the order Vibrionales overwhelmingly dominated the diseased counterparts, except for shrimp 10-BD (Fig. 5b). The order Cytophagales exhibited reduced occurrence in the gut microbiomes of diseased postlarvae compared to the healthy ones. The family Rhodobacteraceae exhibited greater frequency in the gut microbiomes of healthy shrimp postlarvae, whereas the diseased counterparts displayed elevated relative abundances of Vibrionaceae and Pseudoalteromonaceae within the order Vibrionales, except for shrimp 10-BD (Fig. 5c). Pseudoalteromonaceae was not detected in the gut microbiomes of healthy postlarvae. The family Flavobacteriaceae was identified in both healthy and diseased postlarvae’s gut microbiomes. An unassigned genus within the family Rhodobacteraceae emerged as the most abundant in healthy postlarvae’s gut microbiomes and exhibited lower occurrence in the diseased counterparts. The most abundant genera in the gut microbiomes of diseased postlarvae included an unassigned genus within Vibrionales, *Vibrio* sp., and *Pseudoalteromonas* sp. Some genera, such as *Ruegeria*, an unassigned genus within the family Cyclobacteriaceae, *Octadecabacter, Haloferula*, and *Demequia*, displayed modest relative abundances in healthy shrimps (Figs. 5d, 6).

**Figure 4.**
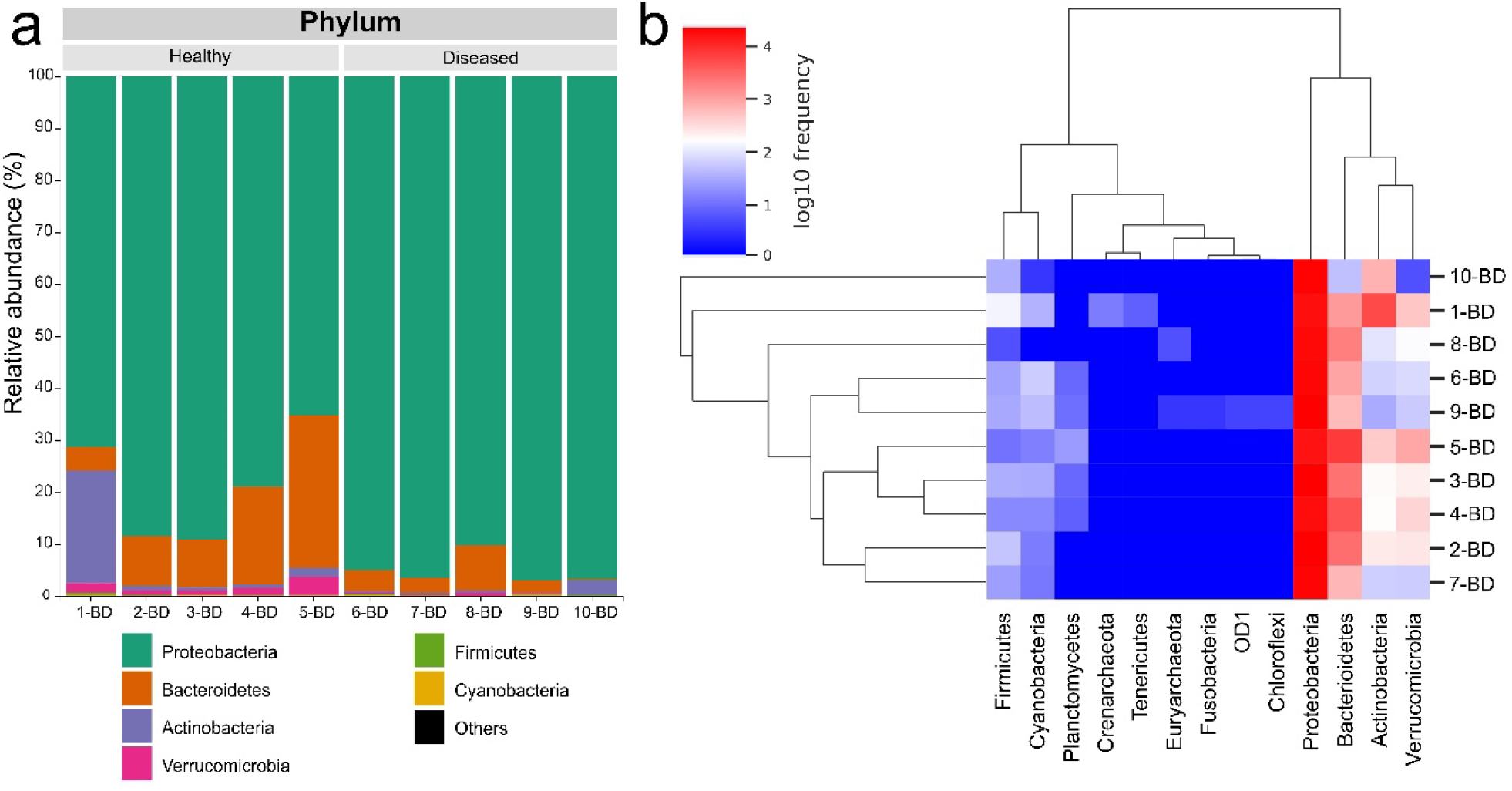
Taxonomic composition at phylum level: (a) barplot and (b) heatmap

**Figure 5.**
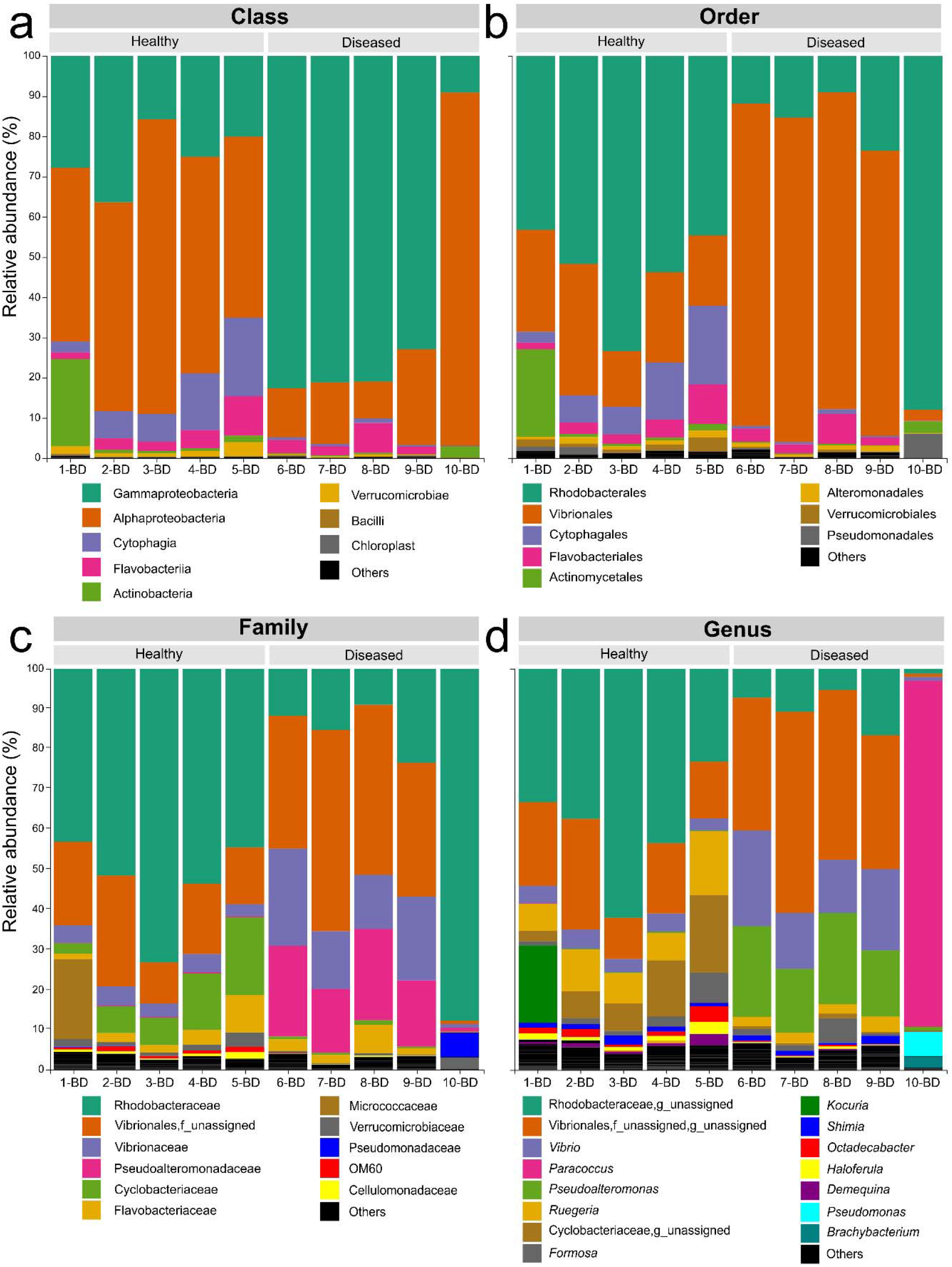
Taxonomic composition barplots indicating relative abundances for each postlarvae’s gut microbiome at (a) class, (b) order, (c) family and (d) genus.

**Figure 6.**
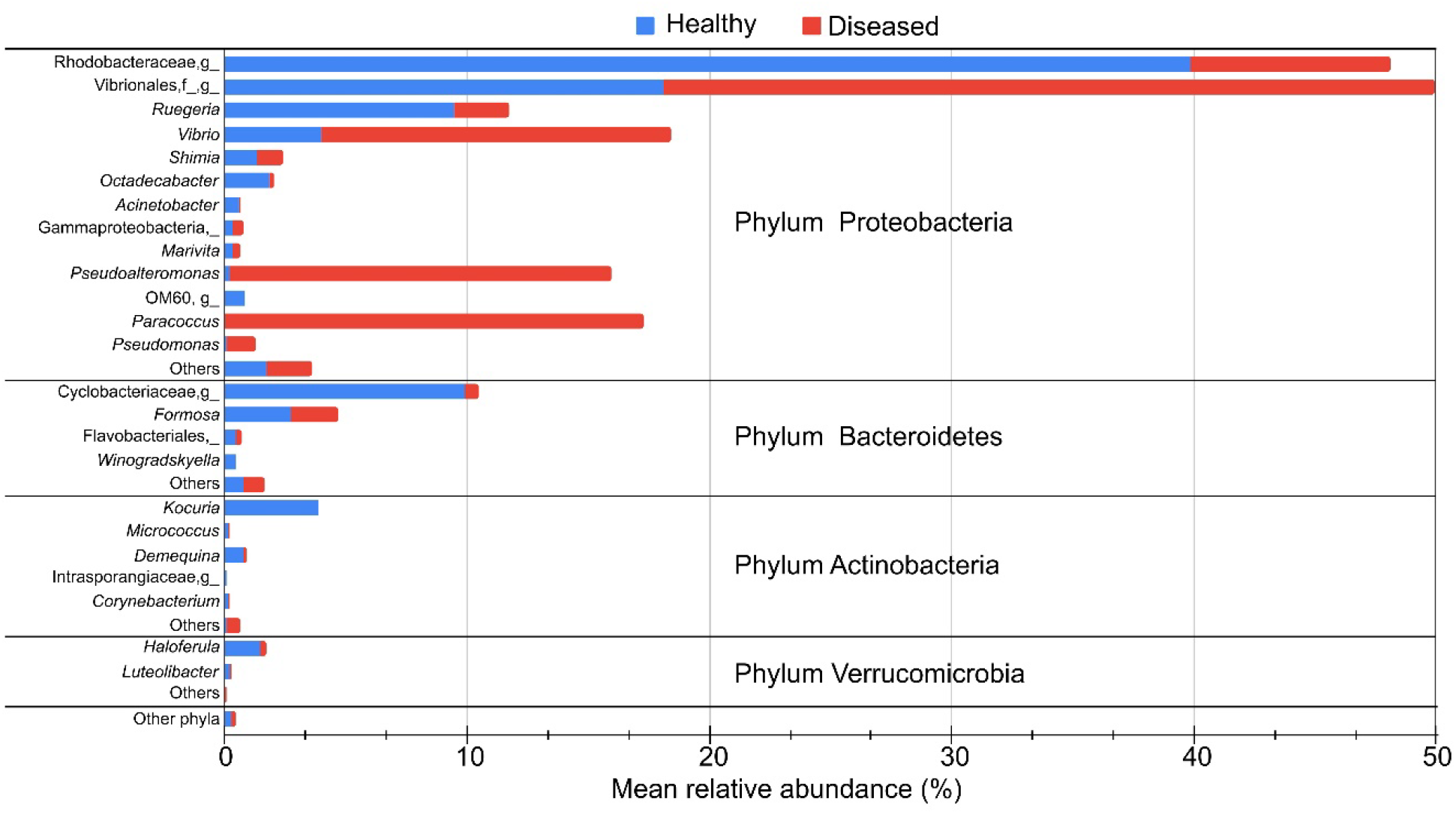
Mean relative abundances as percentages of the most abundant phyla and genera.

## DISCUSSION

In the context of aquaculture, an in-depth comprehension of the gut microbial dynamics is particularly crucial, as it profoundly influences the overall performance and well-being of cultured species. The present study provides insights into the dynamic shifts in diversity and composition observed in the gut microbiome of whiteleg shrimp postlarvae following exposure to AHPND-inducing *V. parahaemolyticus* strain within a closed culture system on the northern coast of Peru.

A highly diverse microbiome is known to exhibit greater stability and resilience in the face of environmental perturbations (Lozupone et al. 2012, Shade 2023). In fact, our findings demonstrate a significant decrease in alpha diversity (Figs. 1a,d) and notable compositional changes (beta-diversity) (Figs. 3a,d) within the gut microbiomes of diseased postlarvae. These results align with previous studies investigating the gut microbiomes of shrimp (Xiong et al. 2015, Chen et al. 2017, Cornejo-Granados et al. 2018, Afrin et al. 2022).

Proteobacteria typically represents the predominant phylum in shrimp gut microbiomes (Holt et al. 2017), in contrast to vertebrates where Bacteroidetes and Firmicutes are more abundant (Rungrassamee et al. 2014, Xiong et al. 2015, 2017, Rungrassamee et al. 2016). Specifically, our study reveals the prevalence of Proteobacteria, Bacteroidetes, Actinobacteria, and Verrucomicrobia, primarily in healthy postlarvae (Fig. 4). These phyla have also been identified in previous investigations involving different life stages (Zhang et al. 2014, Xiong et al. 2015, Huang et al. 2016, Zheng et al. 2016, Dai et al. 2019, Omont et al. 2020). Bacteroidetes, known for their abundance in the marine environment, play a crucial role in the normal gastrointestinal development of aquatic species (Thomas et al. 2011), including shrimps (Cornejo-Granados et al. 2018, Omont et al. 2020). Members of the Actinobacteria phylum are recognized for their production of antibiotic compounds (Huang et al. 2018), however, our study observed lower abundances compared to previous reports (Dai et al. 2018, Landsman et al. 2019).

In the phylum Proteobacteria, the class Alphaproteobacteria dominated the gut microbiomes of healthy postlarvae (Fig. 5a). Members of this class have shown proficient colonization abilities within the healthy gut epithelium, arousing interest in their potential for probiotic production (Xiong et al. 2015, Omont et al. 2020). Conversely, the gut microbiomes of diseased postlarvae exhibited a substantial relative abundance of Gammaproteobacteria, consistent with previous studies (Zhu et al. 2016, Chen et al. 2017, Cornejo-Granados et al. 2018, Afrin et al. 2022). It appears that as the relative abundance of Gammaproteobacteria increases, the diversity of the diseased postlarvae’s gut microbiome decreases (Holt et al. 2020). Interestingly, the class Cytophagia, typically found in rearing water and mysis larvae (Zheng et al. 2017, Wang et al. 2020), was unexpectedly present in both groups.

Orders Rhodobacterales and Vibrionales exhibited the highest abundance in healthy and diseased shrimp postlarvae gut microbiomes, respectively (Fig. 5b). In a previous study investigating the gut microbiomes of whiteleg shrimp postlarvae affected by AHPND, Vibrionales and Mycoplasmatales were identified as the predominant orders (Chen et al. 2017). While Mycoplasmatales are commonly found in the digestive systems of various shrimp species (Huang et al. 2018, 2020), they were not detected in our study.

The family Rhodobacteraceae exhibited predominance in the gut microbiomes of healthy postlarvae (Fig. 5c). Previous studies have reported the presence of this family in the gastrointestinal tracts of healthy shrimp (Huang et al. 2016, Zhu et al. 2016, Chen et al. 2017, Omont et al. 2020). Given their known role in the carbon biogeochemical cycle (Dai et al. 2018) and production of Vitamin B12 (Sañudo-Wilhelmy et al. 2014), they are likely essential components of the normal gut microbiome of whiteleg shrimp (Liu et al. 2019). The family Flavobacteriaceae was detected at variable relative abundances across all samples, except in shrimp 10-BD (Fig. 5c). As no distinct differences were observed between the two groups, it is plausible that members of this family engage in interactions with both commensal and pathogenic species (Kitamoto et al. 2016).

At the genus level, the most prevalent genera in healthy postlarvae’s gut microbiomes were an unassigned genus within Rhodobacteraceae, Ruegeria, an unassigned genus within Cyclobacteriaceae, *Octadecabacter, Haloferula*, and *Demequia*. These genera have previously been identified as members of the healthy shrimp gut microbiome (Zheng et al. 2016, Gainza et al. 2018, Du et al. 2021, Afrin et al. 2022), with the exception of the unassigned genus within Cyclobacteriaceae, which has not been reported before. In the gut microbiomes of diseased postlarvae, the most abundant genera were an unassigned genus within Vibrionales, *Vibrio*, and *Pseudoalteromonas* (Figs. 5d, 6). These genera have been associated with decreased shrimp growth rates (Huang et al. 2010). Several *Vibrio* species, such as *V. harvey, V. parahaemolyticus*, and *V. anguillarum*, are significant pathogens (Huang et al. 2020). Interestingly, *Vibrio* was not the most abundant genus in diseased postlarvae’s gut microbiomes, which aligns with other studies investigating AHPND-affected shrimps (Zheng et al. 2017, Cornejo-Granados et al. 2018, Reyes et al. 2022).

The diseased shrimp 10-BD exhibited a distinct profile compared to the other samples, indicating its outlier status in terms of composition and relative abundance of taxa. Among the most abundant genera identified in this sample were *Paracoccus, Pseudomonas*, and *Brachybacterium. Pseudomonas* has been previously reported in shrimp gut and rearing water (Zhang et al. 2014), however, the presence of *Paracoccus* and *Brachybacterium* as members of the shrimp gut microbiome has not been reported before. This finding suggests a severe dysbiosis that likely led to the displacement of *Vibrio*, although the possibility of handling error or contamination cannot be ruled out.

Deterministic processes primarily govern the bacterial communities in healthy shrimp guts (Xiong et al. 2017). However, in the case of AHPND, the disturbed microbiome appears to be influenced by stochastic processes, along with environmental factors and disease severity (Chen et al. 2017). Inflammatory responses can suppress resident bacteria while favoring the proliferation of opportunistic pathogens like *Vibrio*. Coinfections, secondary infections by opportunistic pathogens, and horizontal transmission of bacteria from the aquatic environment are additional factors to be considered (Carding et al. 2015, Chen et al. 2017, Xiong et al. 2017, Bass et al. 2019, Dai et al. 2019).

Despite the significant changes observed in the composition of diseased postlarvae gut microbiomes reported in this study, further research is necessary to explore additional factors such as geographic location, disease severity, seasonality, and genotype, among others, in order to fully characterize the dysbiosis profile (pathobiome) of an AHPND-affected gut microbiome. The findings presented here contribute to our understanding of the complex ecological interactions within the gastrointestinal tract of shrimps and open avenues for potential therapeutic interventions targeting the gut microbiota to mitigate the impact of AHPND.

## ACKNOWLEDGMENTS

This work received partial support from the Universidad Científica del Sur (UCSUR) Cabieses Grant Nº014-2020-PRE16. We extend our gratitude to CEBAP and Incabiotec S.A.C. for generously providing their facilities and collaborating in shrimp culture, infection experiments, DNA extraction and sequencing. We would like to express our sincere appreciation to César Santos for his invaluable assistance during the experimental phase of this project. Lastly, we are grateful for the assistance provided by Mónica Santa-María and Alonso Reyes Calderón (UTEC) in conducting the initial molecular procedures.

---

**Table Supplementary 1.**
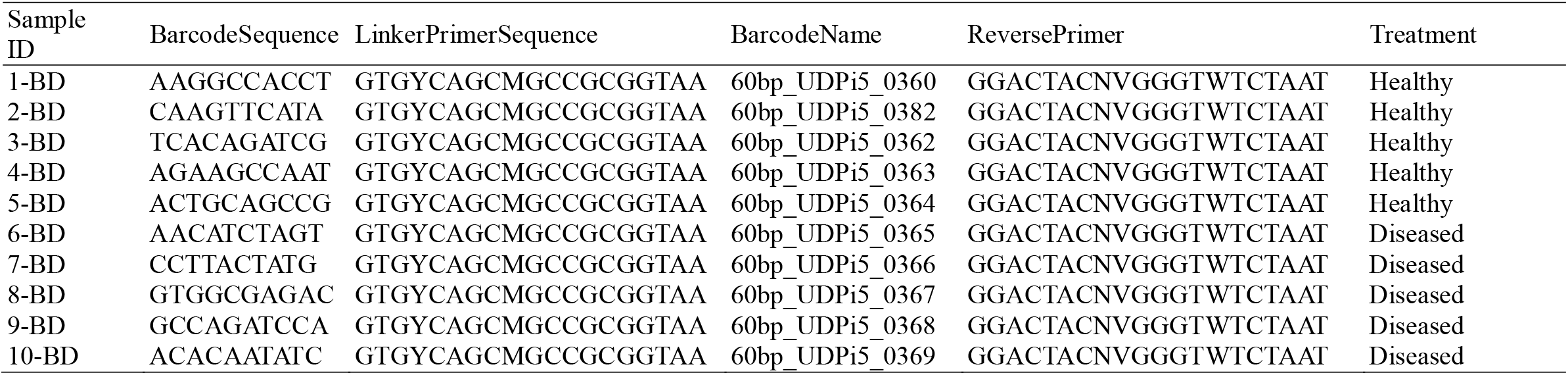
QIIME 2 Metadata showing sample-ids, barcode sequences, primers, and treatments for each analyzed shrimp postlarvae gut microbiome.

**Table Supplementary 2.** Number of sequences before and after of quality control with q2-dada2: filtering, denoising and chimera removal.

| Shrimp<br>(sample-id) | Number of raw<br>sequences<br>(input) | Number of<br>filtered<br>sequences | Percentage of<br>filtered<br>sequences<br>(%) | Number of<br>denoised<br>sequences | Number of<br>non-chimeric<br>sequences | Percentage of<br>non-chimeric<br>sequences<br>(%) |
| --- | --- | --- | --- | --- | --- | --- |
| 1-BD | 34769 | 33669 | 96.84 | 33173 | 27328 | 78.60 |
| 2-BD | 34731 | 33836 | 97.42 | 33542 | 29686 | 85.47 |
| 3-BD | 34753 | 33800 | 97.26 | 33490 | 28719 | 82.64 |
| 4-BD | 34760 | 33447 | 96.22 | 32938 | 26111 | 75.12 |
| 5-BD | 34747 | 33849 | 97.42 | 33350 | 27139 | 78.1 |
| 6-BD | 34755 | 33865 | 97.44 | 33309 | 24177 | 69.56 |
| 7-BD | 34739 | 33904 | 97.6 | 33579 | 24468 | 70.43 |
| 8-BD | 34746 | 33811 | 97.31 | 33301 | 24376 | 70.15 |
| 9-BD | 34749 | 33799 | 97.27 | 33293 | 26379 | 75.91 |
| 10-BD | 34766 | 33868 | 97.42 | 33628 | 25825 | 74.28 |
| Average | 34751.5 | 33784.8 | 97.22 | 33360.3 | 26420.8 | 76.03 |

**Table Supplementary 3.** Alpha diversity measurements of each postlarvae’s gut microbiome and the Mann-Whitney U test significances between treatments (healthy *vs*. diseased) (\**P* < 0.05, ns: non-significant).

| Shrimp | Treatment | ASV richness<br>(S) | Shannon Index<br>(H) | Simpson<br>Index<br>(1-D) | Pielou<br>Index<br>(J) |
| --- | --- | --- | --- | --- | --- |
| 1-BD | Healthy | 94 | 4,03 | 0,89 | 0,62 |
| 2-BD | Healthy | 89 | 3,73 | 0,86 | 0,58 |
| 3-BD | Healthy | 84 | 3,23 | 0,81 | 0,50 |
| 4-BD | Healthy | 84 | 3,69 | 0,87 | 0,58 |
| 5-BD | Healthy | 82 | 4,13 | 0,91 | 0,65 |
| 6-BD | Diseased | 88 | 3,42 | 0,82 | 0,53 |
| 7-BD | Diseased | 53 | 2,71 | 0,71 | 0,47 |
| 8-BD | Diseased | 73 | 3,02 | 0,76 | 0,49 |
| 9-BD | Diseased | 88 | 3,44 | 0,83 | 0,53 |
| 10-BD | Diseased | 33 | 1,80 | 0,61 | 0,36 |
| Mann-Whitney U (*P*-value) |  | 0,2073<sup>ns</sup> | 0,03175<sup>*</sup> | 0,03175<sup>*</sup> | 0,03558<sup>*</sup> |

**Table Supplementary 4.**
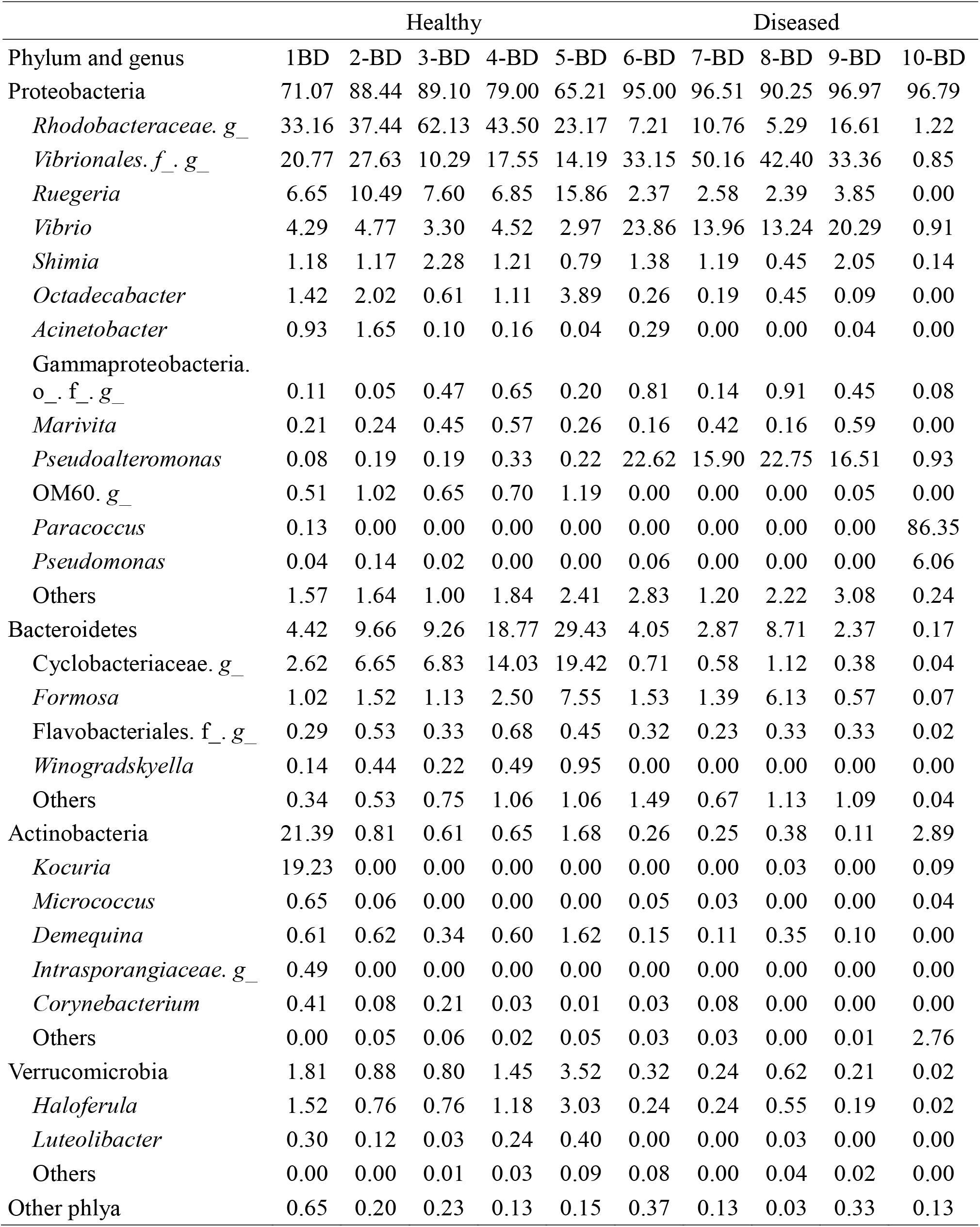
Relative frequencies as percentages (%) of the most abundant phyla and genera per postlarvae’s gut microbiome.

## Notes

### Competing Interest Statement

The authors have declared no competing interest.

